# Blastema cells exhibit intrinsic migratory capacity but fail to induce osteoblast off-bone migration during Zebrafish fin regeneration

**DOI:** 10.64898/2026.08.06.743241

**Authors:** Ivonne M. Sehring, Gilbert Weidinger

**Affiliations:** Institute of Biochemistry and Molecular Biology, Ulm University, Albert-Einstein-Allee 11, 89081 Ulm, Germany

## Abstract

Zebrafish bone regeneration is a highly efficient process, enabling the complete restoration of an amputated fin within few weeks. The hallmark of this epimorphic regeneration is the formation of a blastema atop of a bony fin ray. Osteoblasts near the injury site dedifferentiate and migrate off the bone to contribute to the developing blastema. We show that an injury or a blastema alone is not sufficient to trigger off-bone migration of osteoblasts. Surprisingly, we found that blastema cells themselves possess intrinsic migratory properties. Moreover, when multiple injury sites are present, a preferential distal migration could be observed. We conclude that multiple injuries are hierarchical organized, and that injuries with the highest regenerative potential take priority.

## Introduction

Zebrafish (Danio rerio) exhibit a remarkable capacity for bone regeneration and can fully restore lost appendages, including its bony structures. Thus, the zebrafish caudal fin has become a popular model to study bone regeneration (Gemberling et al., 2013; Pfefferli and Jazwinska, 2015). The bony elements of the fin, the fin rays (lepidotrichia), are segmented, with the segments separated by flexible joints. Each segment is formed by two opposing concave hemirays, which are lined by a single layer of osteoblasts on the inner and outer surface (Figure 1A). The outer layer of osteoblasts adjoins to the basal epidermal layer, while the inner layer faces intraray mesenchyme. Fin growth occurs by distal addition of new segments (Haas, 1962). After fin amputation, a wound epidermis is formed by migration of stump epithelial cells, which rapidly covers the wound (Santos-Ruiz et al., 2002). Regeneration occurs via epimorphic regeneration; that is, a blastema forms atop of each ray. A blastema is composed of a collection of progenitor cells, with intraray fibroblasts giving rise to mesenchymal cells at the core of the blastema (Pfefferli and Jazwinska, 2015; Poss et al., 2003). Osteoblasts close to the amputation plane dedifferentiate, that is they downregulate the expression of mature markers such as *bglap* (Knopf et al., 2011; Sehring et al., 2022; Sousa et al., 2011). This dedifferentiation is negatively regulated by retinoic acid signaling (Blum and Begemann, 2015) and NF-κB signaling (Mishra et al., 2020). Importantly, osteoblasts close to the injury site migrate towards the amputation plane and beyond to contribute to the forming (Geurtzen et al., 2014; Knopf et al., 2011; Sehring et al., 2022). This migratory behavior is specific for osteoblasts after amputation, as osteoblasts show no motility in non-injured fins (Geurtzen et al., 2014), and during addition of new bony segments at the distal tip of growing fins during ontogeny or regeneration, no migration of osteoblasts into a newly forming distal segment is observed (Sehring et al., 2022). Migration of osteoblasts can be analyzed *in vivo* using the transgenic line *bglap:*GFP, a reporter line for mature osteoblasts (Knopf et al., 2011). In mature segments, GFP+ cells can be found predominantly in the center, while they are absent from the segment borders (Figure 1B). Due to the persistence of the GFP protein even if transgene expression is shut down, dedifferentiated osteoblasts can be traced short-therm. After amputation, osteoblasts from segment 0 (the segment through which the amputation is performed) and of the adjacent proximal segment, termed segment -1, migrate towards the amputation plane and beyond into the developing blastema (Figure 1B). In detail, this means that osteoblasts first migrate atop of the bone matrix, and when the amputation plane is reached, they migrate further off the bone matrix (from here one termed “off-bone migration”) into the regenerative mesenchyme.

**Figure 1.**
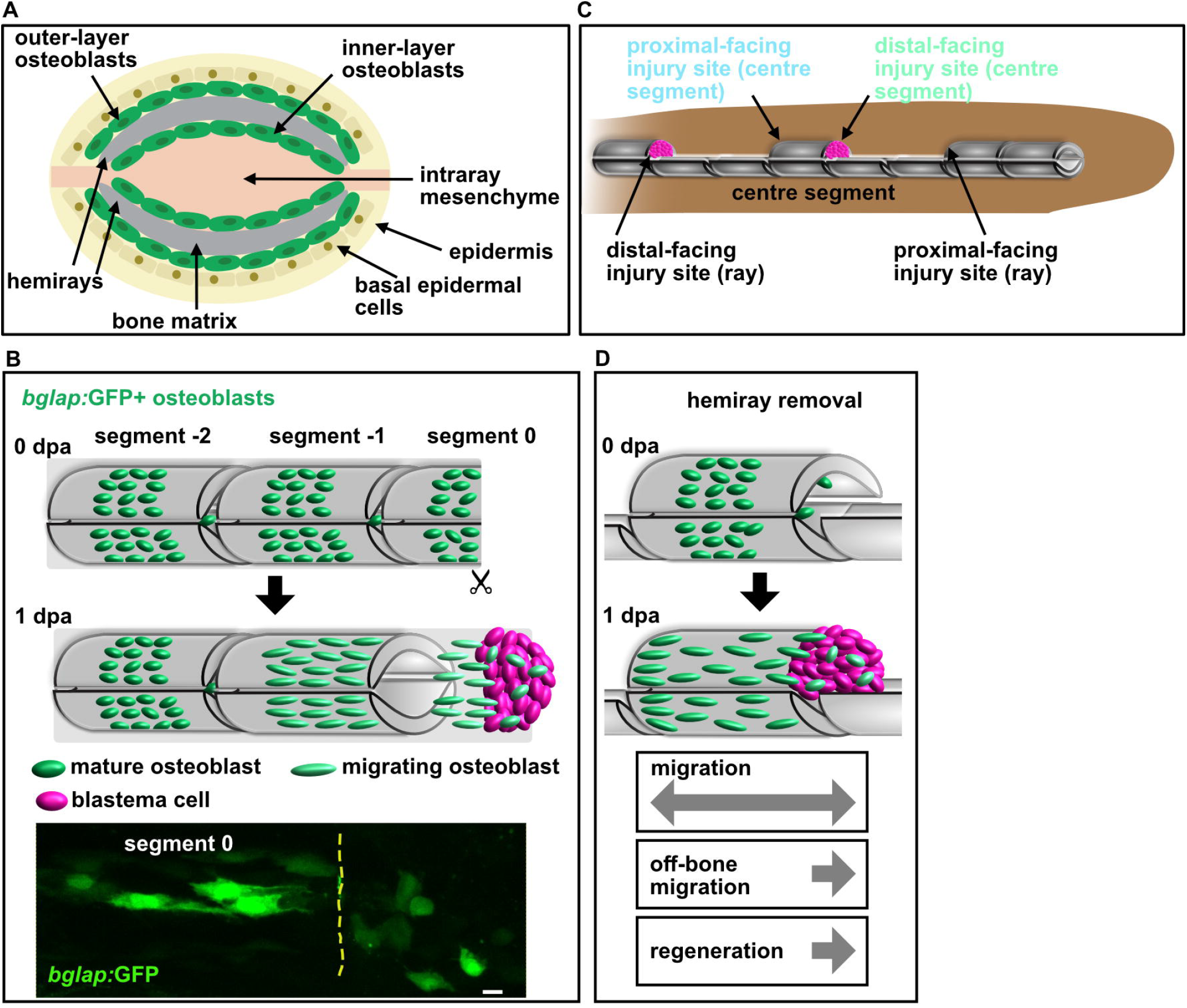
Osteoblast migration and blastema formation in response to fin injury. A) Schematic cross-section of a fin ray. Osteoblasts line the two concave bony hemirays in a single layer. B) Model of osteoblast responses to injury in the zebrafish fin. *Bglap*:GFP+ osteoblasts are located in the center of the mature ray, while they are absent from the segment borders. After fin amputation, osteoblasts migrate towards the amputation plane and contribute to proliferative progenitor cells of the blastema. Microscope image shows migrated *bglap*:GFP+ osteoblasts beyond the amputation plane (indicated by the dashed line). Distal to the right. Scale bar, 10 µm. C) Scheme of the hemiray removal injury model. Removing of hemirays at two locations within one ray results in a proximal-facing and a distal-facing injury site at the centre segment, and additional injury sites in the remaining ray. Blastema only form at the distal-facing injury sites. D) In the hemiray removal injury model, osteoblasts dedifferentiate and migrate at both injury sites, but only distally osteoblasts migrate off the bone, and a blastema is formed. (B, D) are adapted from Sehring et al., 2022.

We have recently established a new injury model where at two locations within one ray, a bone defect is produced, resulting in one intact central bone segment (Figure 1C) (Sehring et al., 2022). In this hemiray injury model (HR2_2 injury), only at the distal-facing injury site a blastema forms and regeneration occurs, eventually leading to polarized bone formation (Figure 1D). In a similar cavity injury model, a consistent outcome could be observed: a blastema formed only at the distal facing injury, and no regenerative growth occurred at the proximal-facing site (Cao et al., 2021). Importantly, in our hemiray removal injury model, both injury sites are equally severed from the stump (and thus cut off from innervation and blood supply), therefore any potential differences in the injury response of the two sites are not due to differences in innervations or blood circulation. Intriguingly however, in our HR2_2 injury, we observed that osteoblasts migration atop of the bone matrix occurred towards both injury sites. Yet only at the distal-facing injury site, they migrated further off the bone matrix into the bone defect (off-bone migration), while at the proximal-facing injury site, they accumulate at the bone ledge (Figure 1D)(Sehring et al., 2022). Since osteoblast migration along the bone matrix occurred towards both injury sites independent of the regenerative outcome, we propose a model in which osteoblast migration atop of the bone is a general response to bone injury. However, our results also indicate that for osteoblasts to migrate off the bone matrix into the bone defect, additional signals are necessary. Blastema formation starts at 12 hour post amputation (hpa) (Poss et al., 2003), while first stray off-bone migrated osteoblasts can be observed at 1 dpa, but are much more frequent at 2 dpa (Knopf et al., 2011; Sehring et al., 2022; Sousa et al., 2011).

We therefore asked if osteoblasts could migrate off the bone – independently off the position of the injury – if a regeneration blastema is present. Blastema formation starts from 12 hours post amputation onwards (Poss et al., 2003; Sehring and Weidinger, 2020), while first migratory osteoblasts (as traced by their *bglap*:GFP expression) can be detected at 1 dpa (Knopf et al., 2011). Therefore, the presence of blastema cells of other lineages might be a required precondition for osteoblasts to migrate beyond the amputation plane. To address this question, we analyzed blastema cell behavior in different HR injury models in combination with various blastema transplantation locations. Our results indicate that the presence of a blastema is insufficient to induce osteoblast migration into the bone defect. Surprisingly, however, our data suggests that the blastema itself has migratory properties and can detect injury signals.

## Results

### A proximal blastema cannot induce osteoblast migration into the bone defect, but migrates distally itself

Xenotransplantation of blastemas is well-established, though previously only transplantations to classical amputation sites have been performed (Shibata et al., 2017; Shibata et al., 2016). To first test whether transplantation of a blastema into a bone defect within a ray would give raise to a new fin ray tissue, we removed hemirays from two adjacent segments within a ray (HR2, Figure 2A), and transplanted a blastema at the distal-facing injury site (mirroring the distal-facing injury site of a classical amputation). To monitor the success of transplantation and survival of the transplanted blastema cells, we used the *ubi*:zebrabow line (expressing RFP in the non-recombined state) as donor. Donor blastemas were generated using the classical fin amputation and harvested at 2 days post amputation (dpa). Repeated imaging over several days showed successful and stable transplantation of the blastema, proliferation of these cells, distal tissue growth, and differentiation into osteoblasts (Figure 2B).

**Figure 2.**
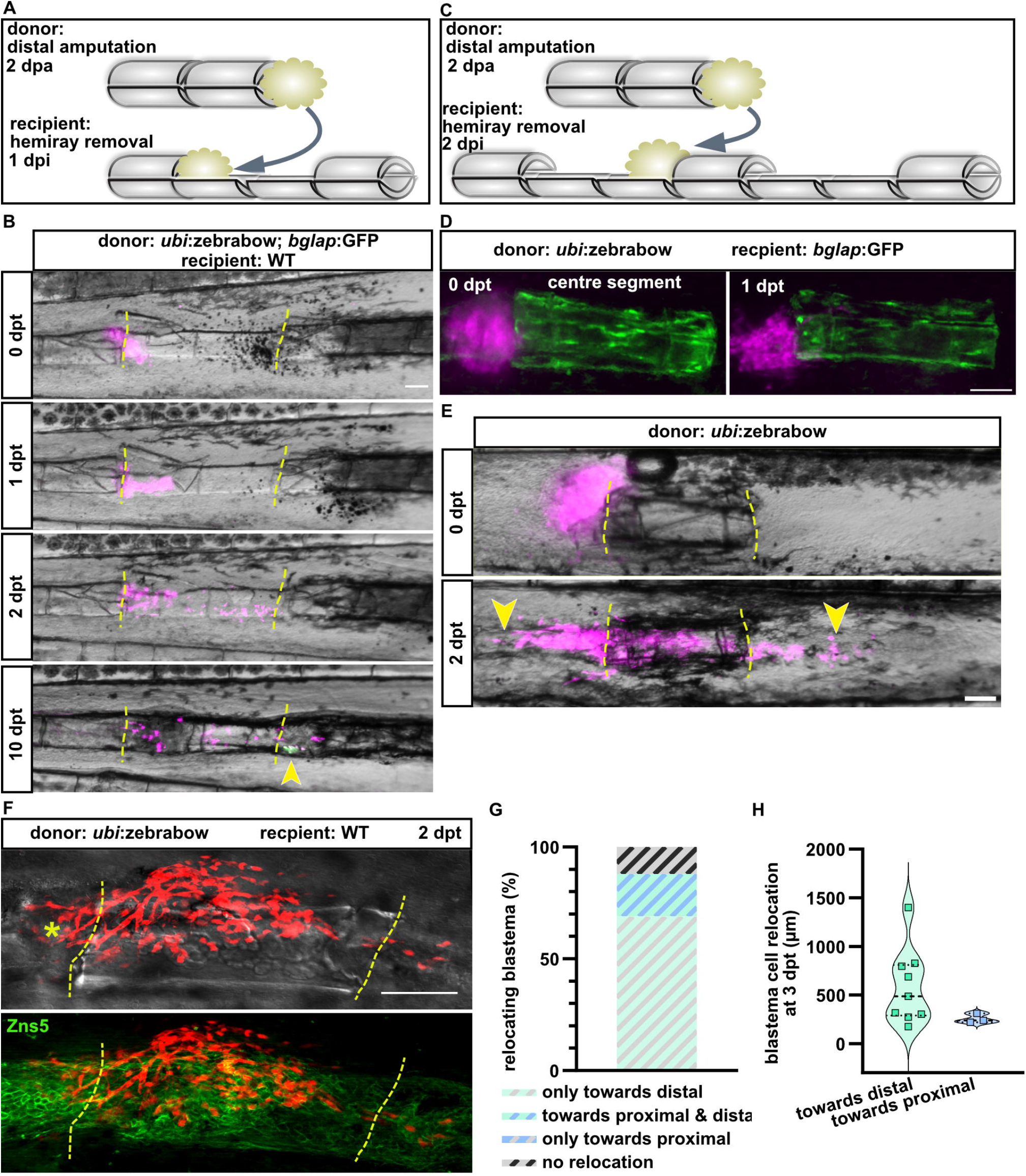
Blastema cells, but not osteoblast relocate in the HR2_2 injury model. A) Scheme of the HR2 injury model and blastema transplantation to the distal-facing injury site of the ray. B) Time-lapse imaging of a HR2 injured ray transplanted with an *ubi*:zebrabow; *bglap*:GFP blastema. At 1 dpt, a distal relocation of blastema cells can be observed. At 10 dpt, expression of *bglap*:GFP can be detected (yellow arrowhead). Dashed lines indicate amputation planes. Distal to the right. Scale bar, 100 µm. C) Scheme of the HR2_2 injury model and blastema transplantation to the proximal-facing injury site of the centre segment. D) Transplantation of a *ubi*:zebrabow blastema to the proximal-facing injury site of *bglap*:GFP fish did not induce osteoblast migration at 1 dpt. Distal to the right. Scale bar, 100 µm. E) Relocation of blastema cells transplanted to the proximal-facing injury site of a centre segment. Dashed lines indicate amputation planes. Yellow arrowheads indicate relocated blastema cells. Distal to the right. Scale bar, 100 µm. F) Co-staining with the pan-osteoblast marker Zns5 highlights the migration of blastema cells atop of the centre segment. Star indicates transplantation site. Dashed yellow lines indicate amputation planes. Distal to the right. Scale bar, 10 µm. G) Distribution of directions of blastema relocations at 2 dpt. n (rays) = 16. H) Absolute maximal length of relocations at 2 dpt. n (towards distal) = 9, n (towards proximal) = 3.

To test whether osteoblast migration off the bone matrix into the bone defect is induced by the developing blastema at the distal-facing injury site of the centre segment, we performed HR2_2 injuries and transplanted blastemas to the proximal-facing injury site of the centre segment (where no endogenous blastema forms; Figure 1C, Figure 2C). To trace osteoblasts in the centre segment as well as to monitor the transplanted blastema cells, we used the *bglap*:GFP osteoblast reporter line as recipient, and the *ubi*:zebrabow line as donor (Figure 2D). The blastemas were transplanted at 2 days post HR2_2 injury (dpi) into the recipient fish to ensure that osteoblasts had already migrated towards the injury sites (as evident by the elongated cell shape of osteoblasts in the centre segment, and their location along the whole segment, Figure 2D). However, at 1 day post transplantation (dpt), no migration of osteoblasts off the bone matrix into the blastema could be observed (n = 5; Figure 2D). Although one day is sufficient to monitor osteoblast migration into a bone defect under standard amputation conditions (Knopf et al., 2011; Sehring et al., 2022), we performed repeated live imaging over several days to ensure that we did not miss any delayed migration. Surprisingly, we instead observed distal-orientated migration of the blastema cells themselves (Figure 2E, F). High-resolution imaging and co-staining with the pan-osteoblast marker Zns5 revealed that blastema cells adopted elongated cell-shapes and migrated atop of the osteoblast along the centre segment (Figure 2F).

### Additional further-distal injury site triggers blastema relocation

Transplantation of a blastema to the proximal-facing injury site of the centre segment in the HR2_2 injury resulted in distal relocation of blastema cells in 69 % of the rays (Figure 2G). While in 19% of cases, both distal and proximal relocation of blastema cells could be observed, no case exhibited only proximal relocation, and only 12% showed no relocation at all. Moreover, the extent distal relocation was significantly more pronounced than proximal relocation (Figure 2H).

Based on these findings, we wondered if a potential signal from the distal injury site could induce the distal blastema relocation. To test this, we returned to the HR2 injury but transplanted blastemas to the proximal-facing injury site (so that no further-distal injury site was present; Figure 3A). In this model, blastema relocation could be observed (Figure 3B), but it was mitigated compared to the HR2_2 model. Only in a few rays (6%), blastema relocation occurred in both directions (Figure 3C), while nearly half of the rays (44%) showed no blastema relocation at all (Figure 3B, C). The number of rays with either proximal or distal relocation were similar (22% and 28%, respectively). Furthermore, the extent of proximal or distal relocation was similar (Figure 3B, D). These results suggest that the presence of a further-distal located injury site influences the proximodistal relocation of blastema cells.

**Figure 3.**
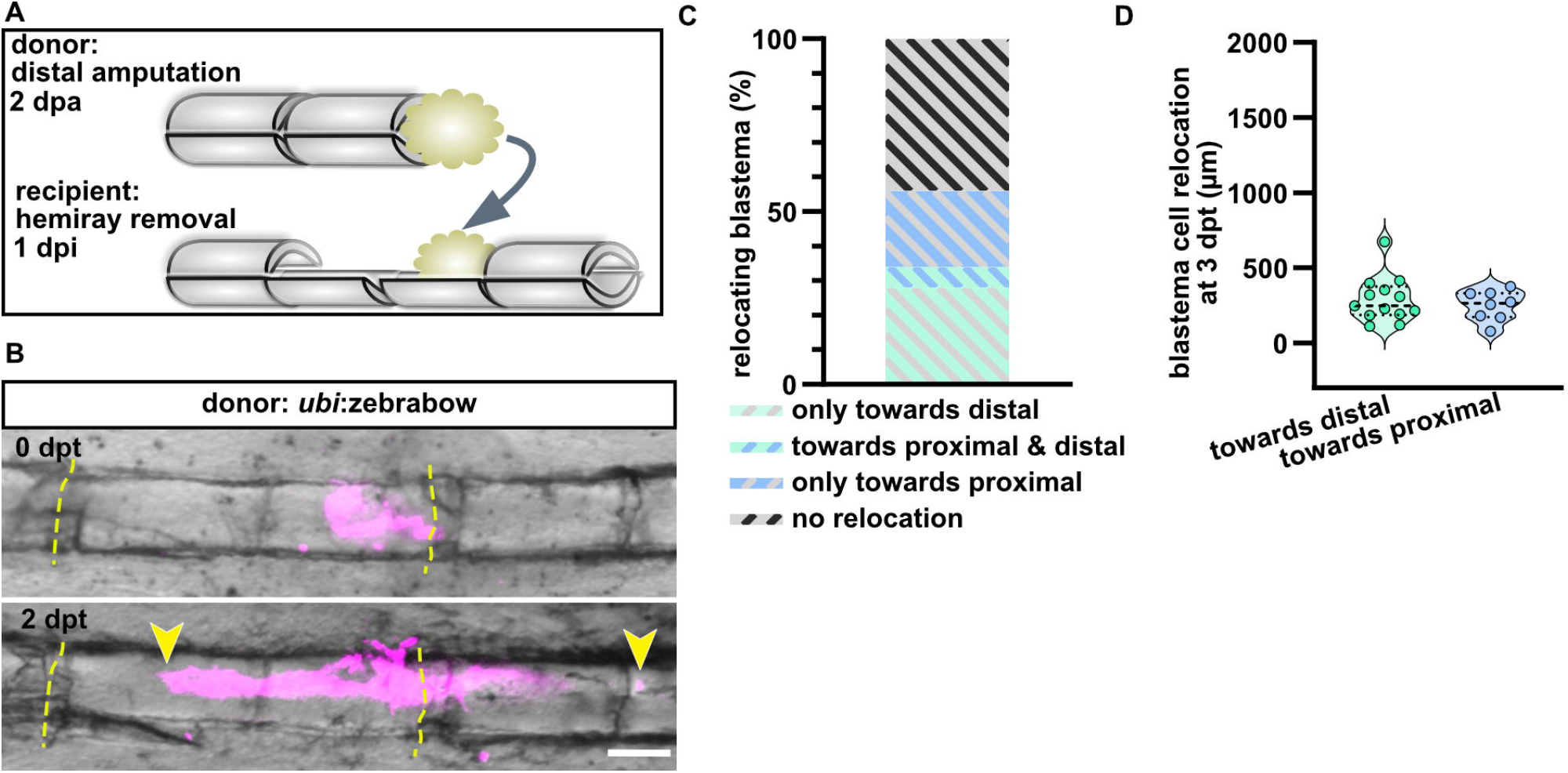
Blastema relocation in the HR2 injury model. A) Scheme of the HR2 injury model and blastema transplantation to the proximal-facing injury site of the ray. B) Relocation of blastema cells transplanted to the proximal-facing injury site of a centre segment. Dashed lines indicate amputation planes. Yellow arrowheads indicate relocated blastema cells. Distal to the right. Scale bar, 100 µm. C) Distribution of directions of blastema relocations at 2 dpt. n (rays) = 18. D) Absolute maximal length of relocations at 2 dpt. n (towards distal) = 13, n (towards proximal) = 8.

To further test this hypothesis, we modified our HR injury model again: instead of one centre segments, two segments remained intact (2HR2_2; Figure 4A). This ensured that the distance between all injury sites was equal (unlike in the HR2_2 model, where the injury sites flanking the centre segments are closer together than to the injury sites of the remaining ray; Figure 2C). Blastema transplantation to the proximal-facing injury site of the centre segment resulted in 44% of rays exhibiting only distal relocation (Figure 4B, C). While 25% displayed both proximal and distal relocation to a similar extent, no cases showed only proximal relocation. 31% showed no relocation, representing a reduction compared to the model with no further-distal injury site (compare Figure 3C and Figure 4C). The extent of proximal or distal relocation was similar (Figure 4D). Together, these data suggest that injury sites can trigger blastema migration, with further-distal injury sites taking priority.

**Figure 4.**
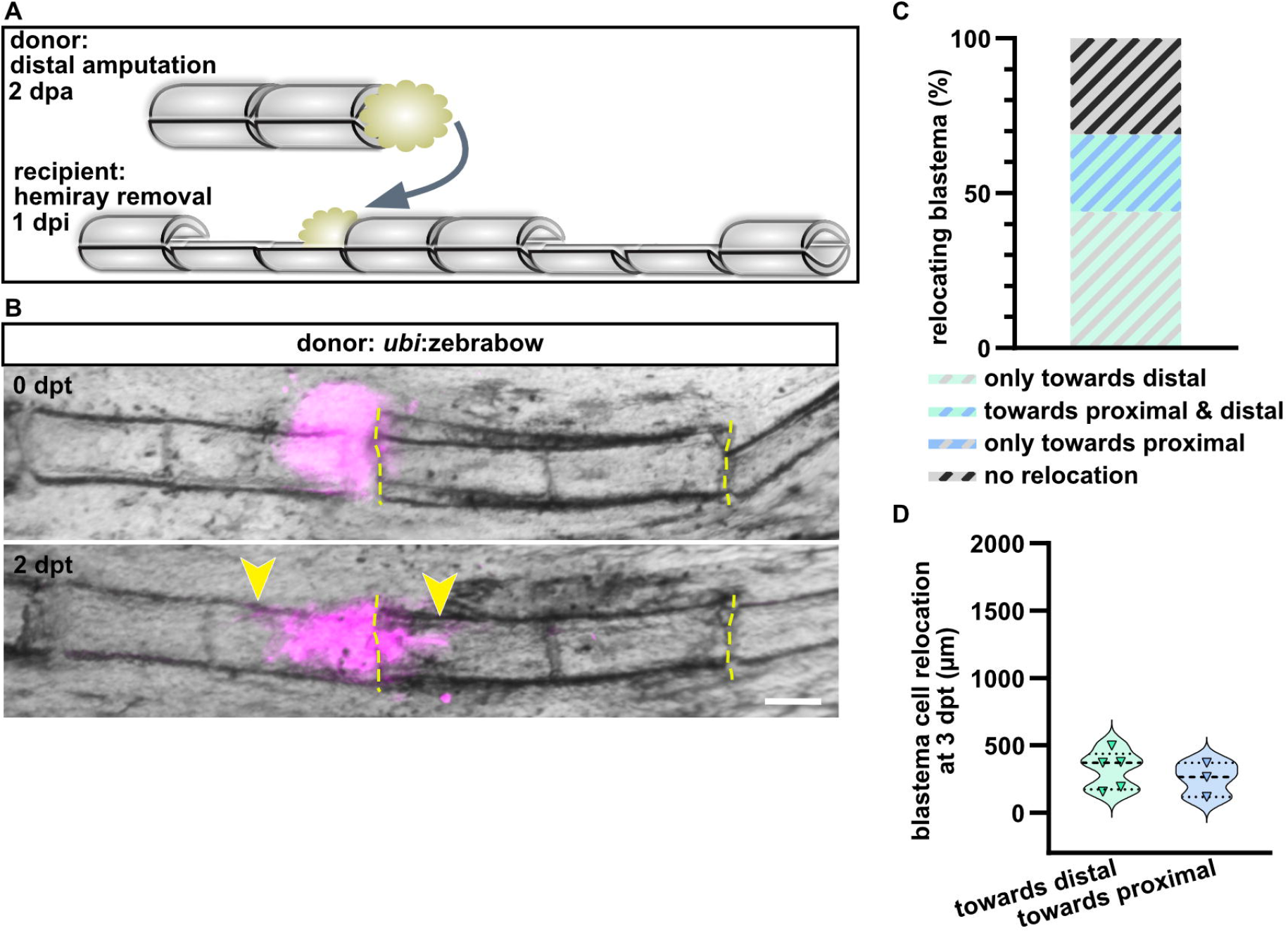
Blastema relocation in the 2HR_2 model. A) Scheme of the 2HR2_2 injury model and blastema transplantation to the proximal-facing injury site of the centre segment. B) Relocation of blastema cells transplanted to the distal-facing injury site of a centre segment. Dashed lines indicate amputation planes. Yellow arrowheads indicate relocated blastema cells. Distal to the right. Scale bar, 100 µm. C) Distribution of directions of blastema relocations at 2 dpt. n (rays) = 16. D) Absolute maximal length of relocations at 2 dpt. n (towards distal) = 5, n (towards proximal) = 3.

### Osteoblast do not migrate into a proximal bone defect

Since the relocation of blastema cell in the HR2_2 injury model hindered our analysis of potential delayed off-bone migration of osteoblasts, we repeated the analysis using the HR2 injury with transplantation to the proximal-facing injury site (Figure 5A). As before, the *bglap*:GFP line was used as recipient to trace osteoblasts, and the *ubi*:zebrabow line as donor to monitor the transplanted blastema cells. While blastema cells persisted at the proximal-facing injury site for four days, no off-bone migration of osteoblasts could be observed (n = 5; Figure 5B). Note that at the distal-facing injury site, osteoblasts can be observed migrating into the bone defect. Our previous data showed that these are indeed migrating osteoblasts and not upregulation of the *bglap*:GFP transgene (Sehring et al., 2022). These data further support our hypothesis that the presence of a blastema is insufficient to trigger off-bone migration of osteoblasts.

**Figure 5.**
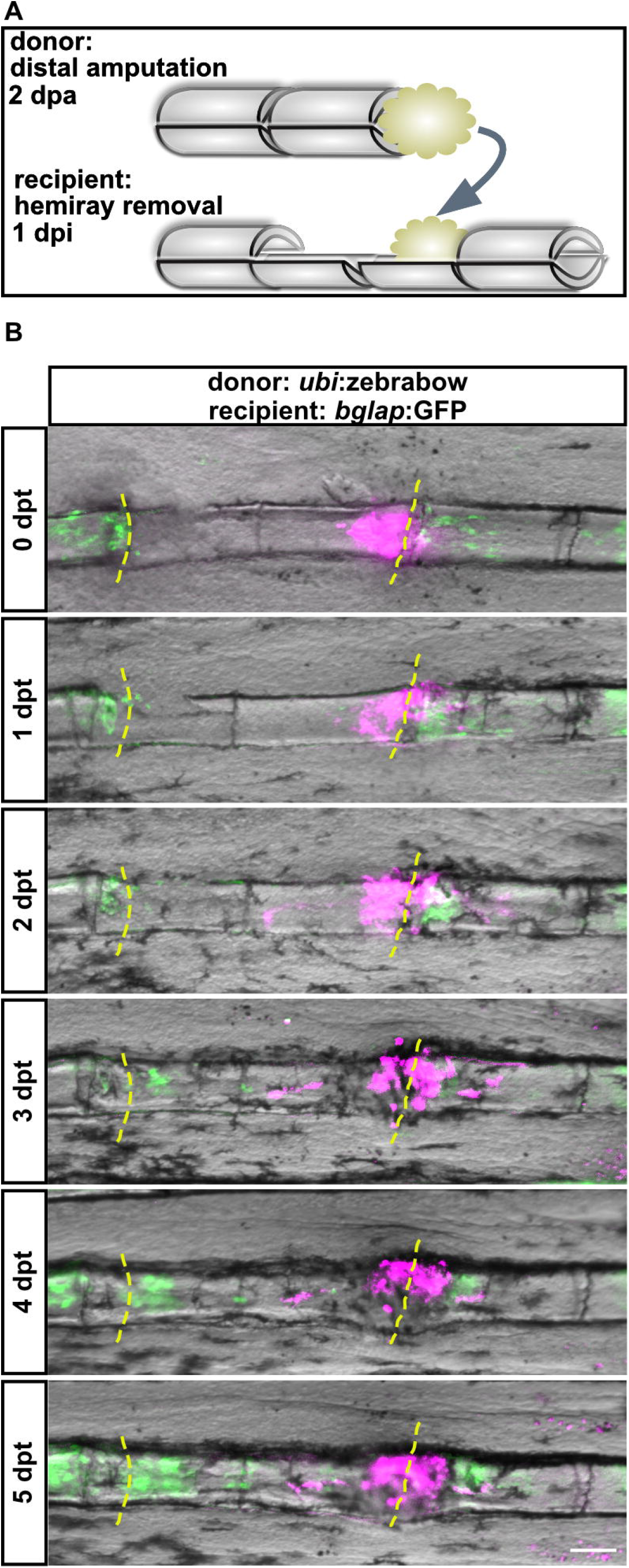
Blastema transplantation does not trigger osteoblast migration. A) Scheme of the HR2 injury model and blastema transplantation to the proximal-facing injury site of the ray. B) Time-lapse imaging of a HR2 injured ray of a *bglap*:GFP transgenic fish transplanted with an *ubi*:zebrabow blastema. No migration of osteoblasts occurred on the proximal-facing injury site, while on the distal-facing injury site, osteoblast can be observed in the bone defect. Dashed lines indicate amputation planes. Distal to the right. Scale bar, 100 µm.

## Discussion

The coordination of various cell types is essential for successful epimorphic regeneration. During zebrafish fin regeneration, the blastema starts to form within 12 hpa through the distal migration of cells from the stump (Poleo et al., 2001; Poss et al., 2003), and is subsequently organized into specific regions with varying proliferation profiles (Nechiporuk and Keating, 2002; Wehner et al., 2014). While the most distal tip of the blastema is thought to hardly proliferate but rather serve as signaling center, the progenitor cells of the proximal blastema proliferate and differentiate, eventually leading to tissue growth and regeneration (Poss et al., 2003). Surprisingly, our data show that blastema cells retain migratory properties even after their initial formation (at 2 dpa), and can respond dynamically to injury signals. Strikingly, our data indicate a preferential distal migration of blastema cells, as transplanted blastemas migrated distally even when initially transplanted to proximal-facing injury sites and thus their migration would lead them away from the bone defect. These observations suggest a hierarchical array of signals dependent on the injury position, which may be critical for guiding regeneration.

A key finding of our work is that the presence of a blastema alone is insufficient to induce off-bone osteoblast migration. This was evident as the transplantation of blastemas to injury sites where no endogenous blastema forms, and no off-bone osteoblast migration occurs, could not trigger migration of osteoblasts off the bone matrix. We have previously shown that dedifferentiation, proliferation and migration atop of the bone matrix of osteoblast in the centre segment occur equally at both injury sites (Sehring et al., 2022). Our new data support our model that migration atop of the bone matrix is a general response to injury, whereas off-bone migration into the bone defect requires additional, as-yet unidentified signals. This distinction is critical, as it implies that these two migrations steps are regulated by different, distinct molecular mechanisms.

In their excavation model, Cao et al. found different calcineurin activity at the two injury sites, and its inhibition could induce blastema formation (Cao et al., 2021). These signals might not be solely generated by the blastema. Potential candidates for such signals include morphogens (e.g., BMPs, Wnt proteins), which are often present as a gradient and could thus contribute to the observed distal preference in cell migration. For example, in the standard amputation model, expression of components of the Wnt/β-catenin pathway is restricted to the regenerative tissue atop of each ray (Wehner et al., 2014); similarly, *bmp4* is likewise exclusively expressed (Murciano et al., 2002). Intriguingly however, in Cao’s excavation model, classical regeneration-associated pathways such as Fgf, Wnt and Notch failed to control polarized regeneration (Cao et al., 2021), suggesting that additional or alternative signals may be involved. Other candidates could be mechanical factors such as tension or compression, which are known to influence cell migration during development and regeneration (Allan and Chaudhuri, 2025). Of note, these signals must differ from those inducing the osteoblast migration atop of the bone matrix, as for the latter, no preferential direction could be observed (Sehring et al., 2022).

The observation that distal injury sites exert a stronger attraction on blastema cells than proximal injury sites raise the question whether there is a spatially organized signal gradient guiding regeneration. However, simple gradient models are not sufficient to explain why, in the HR2_2 model, off-bone migration of osteoblasts can be observed at both distal-facing injury sites, but not at the proximal-facing injury site of the centre segment (which is still distal to the distal-facing injury site of the ray). One possible explanation is that the composition and concentration of signals are not solely dependent on the inherent polarity of the fin (e.g., molecular gradients, (Rabinowitz et al., 2017)) or the absolute proximodistal location of the injury. Instead, they may depend on the direction the injury is facing. Distal-facing injuries might release higher concentrations or distinct combinations of regeneration-promoting factors that direct cell migration. This hypothesis is supported by our 2HR2_2 model, where the distance between all injury sites is evenly distributed, yet blastema cells still preferentially migrated distally. This suggests that blastema cells might be able to distinguish between multiple injury signals, and the biologically most relevant injury – likely the one with the highest regenerative potential – is prioritized.

In summary, this study reveals a previously underappreciated migratory potential of blastemal cells, and underscores the complexity of signaling during regeneration. Both blastema cell migration as well as the off-bone migration of osteoblasts strongly favors distal migration. Therefore, the same signals might be responsible for these migrations. Further work is necessary to identify the specific molecular cues that regulate both blastema cells and osteoblast migrations, and to determine whether hierarchy of signals exists that could regulate the preferred distal migration.

## Methods and Materials

### Animals

All procedures involving animals adhered to EU directive 2010/63/EU on the protection of animals used for scientific purposes, and were approved by the state of Baden-Württemberg (Project numbers 1193 and 1494) and by local animal experiment committees. Fish of both sexes were used. Housing and husbandry followed the recommendations of the Federation of European Laboratory Animal Science Associations (FELASA) and the European Society for Fish Models in Biology and Medicine (EUFishBioMed) (Aleström et al., 2020).

The following pre-existing transgenic lines were used: b*glap*:GFP (Ola.Bglap.1:EGFPhu4008) (Knopf et al., 2011), and *ubi*:zebrabow (Tg(ubb:LOX2272-LOXP-RFP-LOX2272-CFP-LOXP-YFP)) (Pan et al., 2013).

### Fin amputations and hemiray removal

Adult zebrafish were anaesthetized with 625 µM tricaine, and the caudal fins were amputated through the 2nd segment proximal to the bifurcation. For hemiray removal, small surgical blades were used to cut through four joints proximal to the bifurcation in the 2nd ray from ventral and dorsal, and fine tweezers were used to remove the upper hemiray (facing towards the experimenter) of the segments located proximal and distal of one central segment. Fish were allowed to regenerate at 27-28.5°C.

### Blastema transplantation

Blastema transplantation was performed as described in Shibata et al. (Shibata et al., 2017), with the following adaption: An epidermal opening was set at the respective injury site by pushing a thin wolfram wire through the tissue right in front of the centre segment bone. Likewise, the blastema were inserted into the defect using a wolfram wire.

### Whole mount immunohistochemistry

Fins were fixed overnight with 4% PFA at 4°C. After 2 x 5 min washes with PBTx (1x PBS with 0.5% TritonX 100) at RT, fins were transferred to 100% acetone, rinsed once and kept for 3 h at -20°C. Fins were transferred to PBTx, washed 2 x 5 min, 2 x 15 min and incubated for 30 min at RT, followed by blocking in 1% BSA in PBTx for at least 1 h at RT. Primary antibodies were diluted to 1:300 in blocking solution and fins were incubated overnight at 4°C. The next day, fins were washed several times with PBTx and incubated with secondary antibodies at 1:300 dilution in PBTx overnight at 4°C. Fins were mounted in Vectashield (Vectorlabs, H-1000). Primary antibody used in this study weas: mouse anti-Zns5 (Zebrafish International Resource Center, Eugene, OR, USA, RRID:AB_10013796).

### Imaging

Images were acquired with a Leica M205FA stereo microscope and display live fluorescence of fluorescent proteins. Fish were anaesthetized with 625 µM tricaine, and placed on object slides. After imaging, fish were returned to the system at 27-28.5°C. High resolution optical sections of whole-mount antibody staining were obtained with a Zeiss AxioObserver 7 equipped with an Apotome and processed using Fiji (Schindelin et al., 2012).

## Acknowledgements

We thank Doris Weber and Janet Köhler for their contributions to fish care. This work was funded by the Deutsche Forschungsgemeinschaft (DFG, German Research Foundation) – Project-ID 251293561 – SFB 1149.

## Data Availability

The datasets generated during and/or analysed during the current study are available from the corresponding author on reasonable request.

